# In-host evolution of classic to convergent *Klebsiella pneumoniae* sequence type 147 isolates and impact of associated capsular changes on different morphotypes

**DOI:** 10.1101/2024.02.13.580130

**Authors:** Katharina Sydow, Eyüp Doğan, Michael Schwabe, Stefan E. Heiden, Muhammad Moman Khan, Justus U. Müller, Jürgen A. Bohnert, Daniel Baecker, Rabea Schlüter, Peter Schierack, Elias Eger, Evgeny A. Idelevich, Karsten Becker, Katharina Schaufler

**Author notes:** Corresponding author; Helmholtz Institute for One Health, c/o University Medicine Greifswald, Epidemiology and Ecology of Antimicrobial Resistance, Fleischmannstraße 42, 17489 Greifswald, Germany. These authors contributed equally (Karsten Becker, Katharina Schaufler). **Abbreviations** g/d grey and dry; referring to the colony morphotype on blood agar.

## Abstract

*Klebsiella pneumoniae*, an important opportunistic pathogen, has long been categorized into two distinct pathotypes: the often multidrug-resistant classic (cKp) and the highly virulent hypervirulent (hvKp). However, a recent global trend has witnessed the emergence of convergent strains, seamlessly combining antimicrobial resistance with hypervirulence. Our study delved into a series of *K. pneumoniae* isolates sourced from the same patient, all belonging to the international, high-risk clonal lineage of sequence type 147. As reported in a previous study, these isolates exhibited diverse morphotypes on blood agar, ranging from small white to normal-sized white, grey, or grey and dry (g/d) colonies.

Through an interplay of omics and phenotypic experiments, we unraveled the intricate mechanisms governing these distinct colony morphologies and their implications on bacterial virulence and resilience. While the earlier isolates demonstrated modest levels of resistance and virulence, their later counterparts showed significantly heightened levels, attributed to the acquisition of additional plasmids. Bioinformatics analysis unveiled a chromosomal insertion of a hybrid plasmid in one isolate, marking an unprecedented in-host microevolution from the classic to the convergent pathotype.

All morphotypes exhibited positive insertion sequences around or within the K loci, with the grey or g/d phenotypes arising from impaired K loci. Despite lower serum resistance, these morphotypes demonstrated superior adhesion to human epithelial cells. Interestingly, while capsule-deficient strains are conventionally associated with decreased virulence, our isolates displayed high mortality rates in the *Galleria mellonella* infection model.

In conclusion, our findings not only provide unprecedented insights into in-host microevolution within a patient, transitioning from the classic to the convergent pathotype, but also contribute significantly to the understanding of the diverse morphotypes exhibited by *K. pneumoniae*.

## 1. Introduction

*Klebsiella pneumoniae* is a natural part of the human gut microbiome, but as an opportunistic pathogen, also associated with several infections [1]. Traditionally, it has been differentiated into classic (cKp) and hypervirulent (hvKp) pathotypes [1]. While cKp frequently, but not always, carries multiple antibiotic resistances, it mainly causes infections, such as pneumonia, urinary tract infections, and bacteremia, in immunocompromised patients [1]. In contrast, hvKp representatives are mostly antibiotic-susceptible, but are capable of causing community-acquired infections, which are characterized by especially severe courses of infection due to their enhanced ability to metastasize [1]. Even more worrying, in recent years, a convergent pathotype has emerged, which combines hypervirulence with multidrug resistance traits [2–5].

Usually, all *K. pneumoniae* pathotypes produce virulence-associated capsular polysaccharides, but hvKp strains tend to produce higher amounts and are often associated with hypermucoviscosity (HMV) [6]. It is also known that mutations in capsule biosynthesis pathways impact capsule composition, resulting in hyper-capsules or capsule deficiencies [7]. Together with other features, *rmpA* regulates the mucoid phenotype and is mostly plasmid-encoded [6,8,9], sometimes even on so-called hybrid or mosaic plasmids that harbor both virulence and antimicrobial resistance determinants [2]. In addition, such mobile genetic elements can be integrated into the chromosome, which results in stable “fixation” of the respective beneficial genes [10].

Recently, we described *K. pneumoniae* isolates belonging to the international, high-risk clonal lineage of sequence type (ST)147 that showed different colony morphologies on blood agar (Tab. 1) [11]. The isolates were obtained at five different time points from urine and tissue of a patient with shrapnel shell injury. We identified four distinct phenotypes/morphologies for the isolates from the last two time-points, i.e., presenting as (i) white/whitish and (ii) grey colonies, respectively, with a smooth and shiny surface, (iii) grey and dry colonies (g/d), and (iv) small colony variants (SCVs), which were white and smooth [11]. When sub-cultivated on agar plates, the small colonies steadily split again into the four different morphotypes, a phenomenon that we subsequently termed “hyper-splitting” (Fig. 1) [11]. Antibiotic susceptibility testing (AST) revealed varying minimum inhibitory concentrations for isolates from the different sampling time points, which was also dependent on the respective phenotype [11].

**Figure 1.**
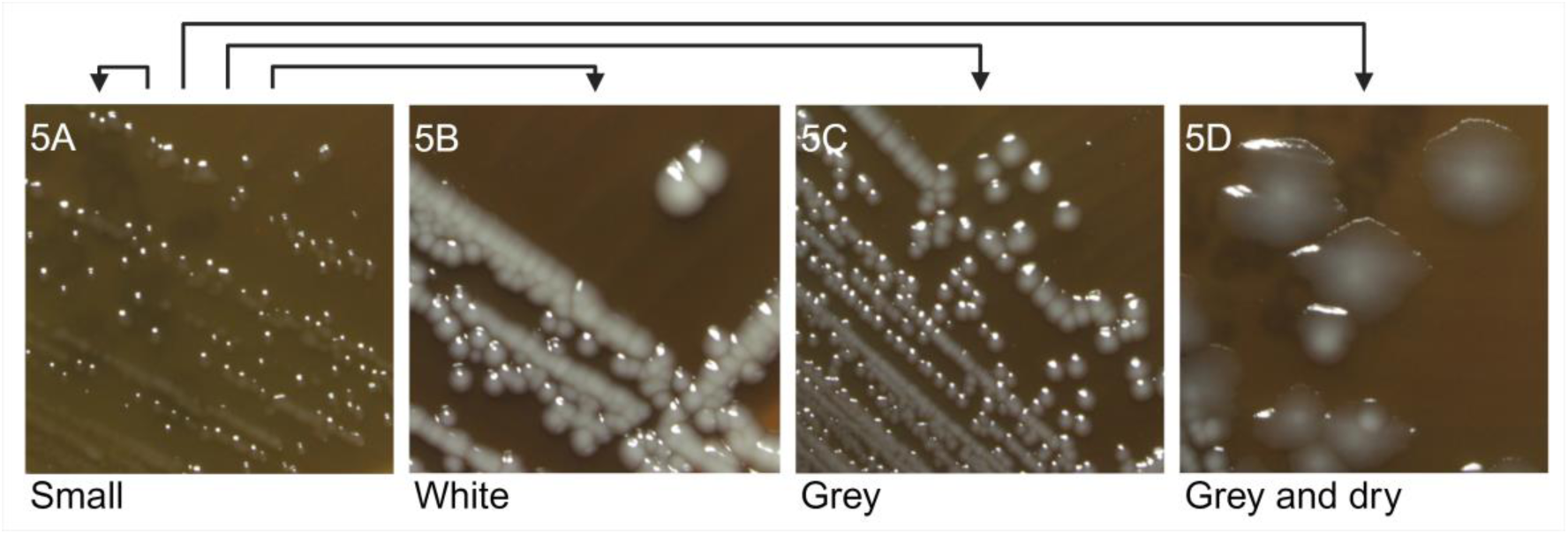
Phenotypic hyper-splitting of the small phenotype into distinct phenotypes; representative morphologies shown for isolates 5A-D, obtained from the last sampling time point, and grown on blood agar.

**Table 1:**
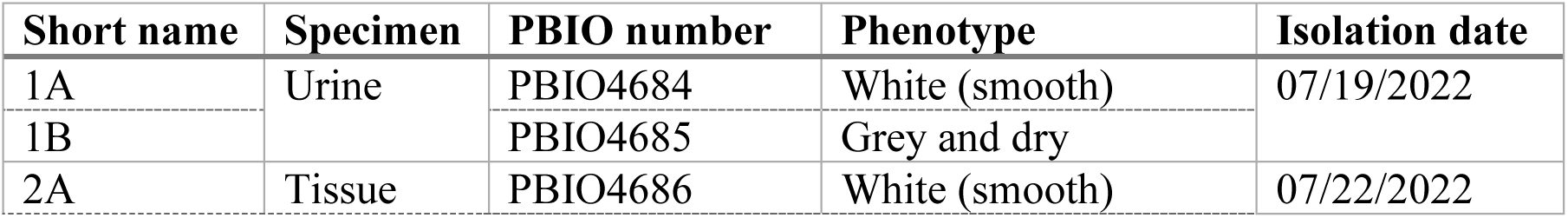
Overview of *K. pneumoniae* isolates with different phenotypes.

*K. pneumoniae* isolates displaying the typical (“normal”) morphology usually present dome-shaped, white or whitish, smooth and shiny colonies on blood agar, which is mostly independent of the pathotype. However, HMV can be occasionally observed as positive string test and mucoid morphologies. While SCVs have been frequently described in staphylococcal species [12,13] and, to a lesser extent, in some Gram-negative bacteria such as *Escherichia coli* and *Pseudomonas aeruginosa* [14–16], only limited variation has been described for *K. pneumoniae*, e.g., after exposure to antibiotic compounds [17] or phages [18]. Morphological changes range from single cells with different appearance [18] to colonies with altered surfaces [7,19]. Notably, these alterations all relate to changes in the capsule, which affects antibiotic susceptibility [19], biofilm formation [7] or virulence [18]. In another previous study, we have described a *K. pneumoniae* isolate with sponge-like colonies on blood agar that produced unusually high amounts of biofilm-related cellulose and contained insertion sequences within the K and O loci [20].

Here, beyond the superficial description [11], we characterized the underlying mechanisms of the morphotypes in detail and tracked the in-host evolution of these isolates. By combining genomics and transcriptomics with phenotyping, we also explored the hyper-splitting phenomenon and revealed associations between the phenotypes and virulence of this important international, high-risk clonal *K. pneumoniae* lineage.

## 2. Materials and Methods

### 2.1. Bacterial samples

*K. pneumoniae* isolates that showed up to four different phenotypes were obtained in a previous study (Tab. 1; [11]). We stored them at -80 °C in phosphate buffered saline (PBS) with 20 % (*v/v*) glycerol (anhydrous; Merck, Darmstadt, Germany) and cultivated the isolates on Columbia blood agar plates containing 5 % sheep blood (Mast Diagnostica, Reinfeld, Germany).

For phenotypic assays, additional strains served as reference and control (Tab. 2), including a convergent type *K. pneumoniae* ST307 from a clonal outbreak at the same university hospital (PBIO1953, [3]), a hypervirulent *K. pneumoniae* ST420 (PBIO2030, [21]), a hypervirulent *K. pneumoniae* ST86 isolate (hvKP1, [22,23]), an *E. coli* K12 (ST10) (W3110, [24]) and, for capsule staining, the quality-control strain *K. pneumoniae* ATCC 13883.

**Table 2:**
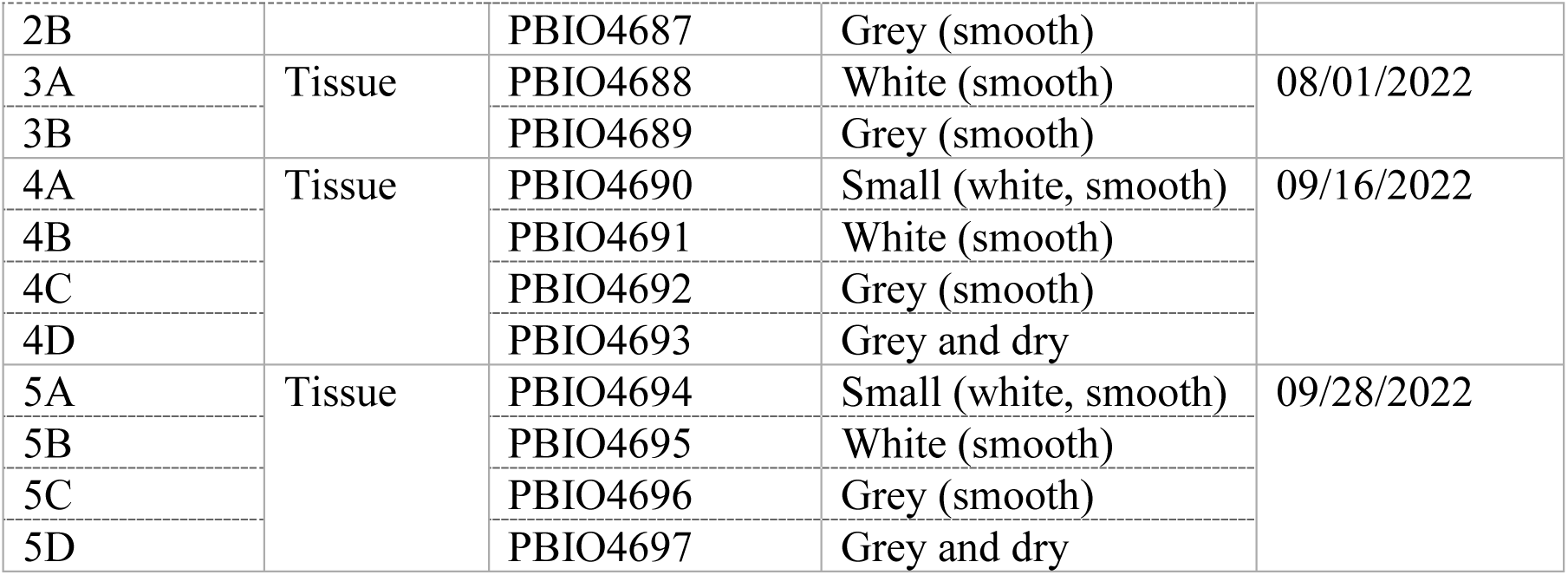
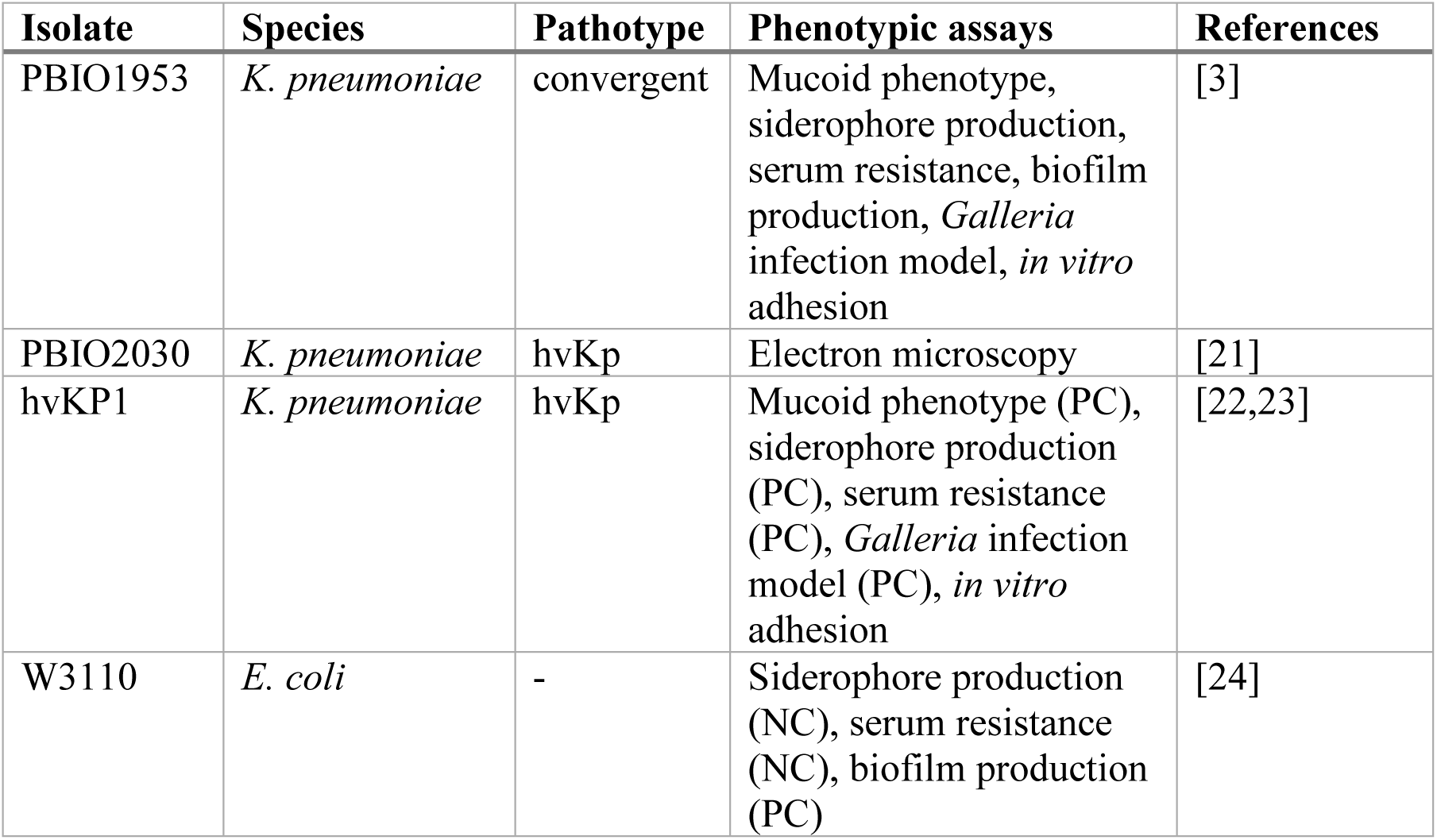
Isolates used as reference and control for phenotypic assays. PC: positive control, NC: negative control.

### 2.2. Genomic analyses

While short read data of all 14 isolates were obtained in the previous study [11], we now performed hybrid sequencing (Illumina and Nanopore) for isolates 5A-D. DNA was extracted using the MasterPure DNA Purification kit for Blood, v.2 (Lucigen, Middleton, WI, USA) according to the manufacturer’s instructions and quantified with the Qubit 4 fluorometer and the dsDNA HS Assay kit (Thermo Fisher Scientific, Waltham, MA,

USA). Samples were sent to the Helmholtz Center for Infection Research (Braunschweig, Germany) for Illumina short-read (NovaSeq 6000, 2 × 150 bp reads) and Oxford NanoPore long-read (FLO-MN106D, R9.4.1, MinION platform) sequencing. Illumina short-reads were investigated with FastQC v.0.11.9 (https://www.bioinformatics.babraham.ac.uk/projects/fastqc/) and multiQC v.1.14 [25] prior and after quality and adapter trimming with fastp v.0.23.2 [26,27], and used together with the Nanopore long-reads as input for Unicycler v.0.5.0 [28]. Resulting genomes were visualized and inspected in Bandage v.0.8.1 [29]. Annotation was performed with bakta v.1.7.0 and the corresponding bakta database v.5.0 [30]. Using the short-read data previously obtained for all 14 isolates [11], potential mutations were investigated with breseq v.0.38.1 [31] using the closed genome sequences of isolate 5B as reference and the trimmed short-reads as input. Results were manually investigated and compared to each other.

To examine the gene expression of the four isolates (5A-D), RNA sequencing and a differential gene expression analysis was performed. Therefore, isolates were grown on blood agar plates and incubated at 37 °C overnight. Total RNA was extracted with the RNeasy kit (Qiagen, Hilden, Germany). Additionally, RNA samples were purified with the rDNase set and the RNA Clean-up kit (Macherey-Nagel, Düren, Germany). RNA quality was checked with the Prokaryote Total RNA Nano Series II chip using the Agilent Bioanalyzer 2100 (Agilent Technologies, Santa Clara, CA, USA) according to the manufacturer’s instructions, and samples were sent to Novogene (Cambridge, UK) for sequencing (NovaSeq 6000 S4, paired end [2 × 150 bp reads], rRNA-depleted, stranded). RNA short-reads were adapter and quality trimmed with Trim Galore v.0.6.8 (https://www.bioinformatics.babraham.ac.uk/projects/trim_galore/) and investigated with FastQC and multiQC prior and after trimming. Trimmed reads of all four samples were mapped with bowtie2 v.2.5.1 [32] to the closed assembly of isolate 5B and formatted with samtools v.1.13 [33]. FeatureCounts v.2.0.1 (stranded-mode) [34] was used to count read-mappings on coding features. The final differential gene expression analysis was performed in R v.4.3.1 (https://www.r-project.org/) with DESeq2 v.1.40.0 [35]. Genes were considered as differentially expressed when the absolute log_2_-fold change was greater or equal to 2 and the adjusted *p* value less than 0.05.

To confirm possible chromosomal plasmid integration, custom primers (primer pair [PP]1 and 2) in combination with primers for the housekeeping gene *infB* were used (Tab. 3; Eurofins GSC Lux, Luxembourg, Luxembourg).

**Table 3:**
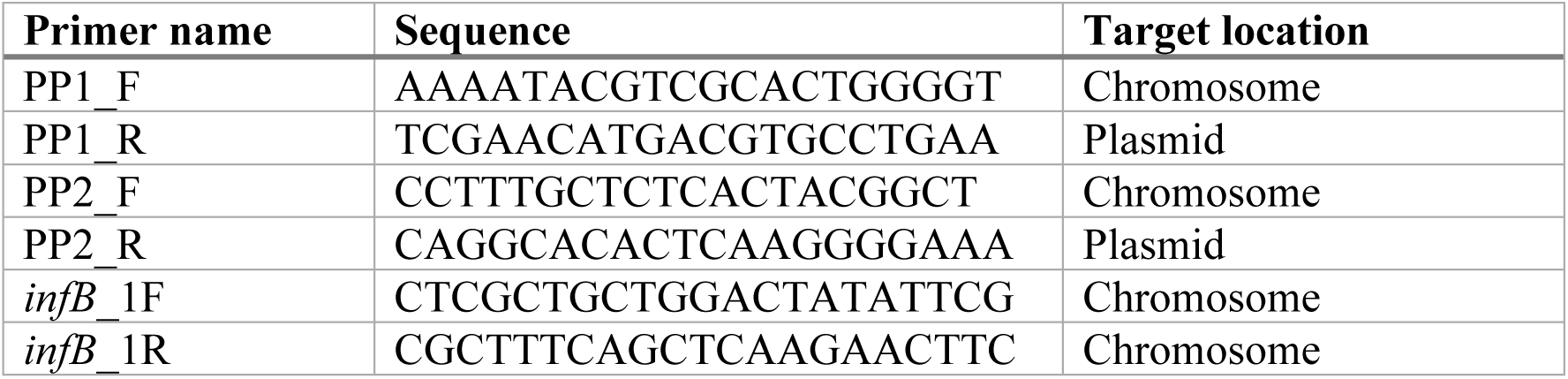
Primers used for confirmation of a chromosomally integrated plasmid.

For each sample, 12.5 µL DreamTaq Green PCR master mix (Thermo Fisher Scientific, Waltham, MA, USA), 1 µL of each primer, 8.5 µL nuclease-free water (Thermo Fisher Scientific, Waltham, MA, USA) and 2 µL DNA template were used. PCR products were amplified with a peqSTAR thermocycler (VWR, Radnor, PA, USA) and the following conditions: initial denaturation for 5 min at 95 °C, 30 cycles of denaturation for 30 s at 95 °C, annealing for 30 s at 58 °C (for PP1 and PP2) or 53 °C (for *infB*) and elongation for 4 min 30 s (for PP1 and PP2) or 1 min (for PP1_F with PP2_R and for *infB*) at 72 °C, ended by 10 min at 72 °C. Samples were separated on a 1.2 % agarose gel stained with 0.01 % (*v/v*) GelGreen Nucleid Acid Stain (Merck, Darmstadt, Germany) in TBE buffer for 2 h at 120 V; GeneRuler 1 kb Plus DNA ladder (Thermo Fisher Scientific, Waltham, WA, USA) was used as a size marker. Results were documented with Quantum CX5 and BioVision v.17.06 software (Vilber, Collégien, France).

### 2.3. Phenotypic assays

#### 2.3.1. Mucoid phenotype

Hypermucoviscosity was determined using the string test. Here, a sterile inoculation loop was used to stretch single colonies on an agar plate. In case a string of 5 mm or longer formed, the test was defined positive [36]. Additionally, HMV was assessed with a sedimentation assay as described previously [21]. Briefly, a bacterial suspension with 0.5 McFarland standard turbidity was prepared in 0.9 % (*w/v*) NaCl, 50 µL of this suspension were added to 5 mL lysogeny broth (LB; Carl Roth, Karlsruhe, Germany) and incubated under shaking conditions (130 rpm) at 37 °C for 24 h. Then, 1.5 mL were centrifuged in 2 mL reaction tubes (Carl Roth, Karlsruhe, Germany) at 1,000 × *g* and room temperature (RT) for 5 min. From the supernatant, 200 µL were added to a 96-well plate (Nunc, Thermo Fisher Scientific, Waltham, WA, USA), 200 µL of the initial culture were transferred separately and the optical density was measured at *λ* = 600 nm (OD_600_) using a microplate reader (CLARIOstar Plus, BMG LABTECH, Ortenberg, Germany).

#### 2.3.2. Biofilm production

Biofilm production was determined with a crystal-violet (CV) assay as described previously [24]. Overnight cultures in LB were diluted 1:100 in M9 medium (MP Biomedicals, Irvine, CA, USA) supplemented with 1 mM magnesium sulfate (Carl Roth, Karlsruhe, Germany) and 0.4 % glucose (Carl Roth, Karlsruhe, Germany) and incubated under shaking conditions (130 rpm) at 37 °C until 0.5 McFarland standard turbidity was reached. Bacterial suspensions were 1:10 diluted and 200 µL were transferred into a 96-well plate. Triplicates of sterile medium were used as control and blank. After incubation at 28 °C for 24 h, OD_600_ was measured with the microplate reader before the 96-well plate was washed thrice with deionized water and air-dried for 10 min. Fixation of cells with 250 µL methanol (Merck, Darmstadt, Germany) for 15 min and subsequent air-drying for 10 min followed. Then, cells were stained with 250 µL of a 0.1 % (*w/v*) aqueous CV solution (Sigma-Aldrich. St. Louis, MO, USA) for 30 min, washed thrice with deionized water and air-dried for 10 min. Bound CV was dissolved in 300 µL of a 80 % ethanol (99.8 % (*v/v*); Carl Roth, Karlsruhe, Germany) and 20 % acetone (Merck, Darmstadt, Germany) solution with horizontal shaking (200 rpm) at RT for 30 min. Lastly, 125 µL of the solution were transferred to a new 96-well plate and OD was measured at *λ* = 570 nm using the microplate reader. Specific biofilm formation (SBF) was calculated according to the following formula [37]: SBF = (B - NC)/G, with B as the OD_570_ of the stained bacteria, NC as the OD_570_ of the stained control wells to eliminate the fraction of CV adhering to the polystyrene surface due to abiotic factors, and G is the OD_600_ representing the density of cells grown in the media.

#### 2.3.3. Serum resistance

In order to test the survival in human blood, serum resistance was tested in 50 % human serum as described previously [21] with slight variations. Instead of using overnight cultures, colonies were taken directly from blood agar plates. The colonies were suspended in 1 mL PBS until the OD_600_ reached 0.5 McFarland standard turbidity. On a 96-well plate, the bacterial suspension and human serum (Merck, Darmstadt, Germany) were mixed in a 1:1 ratio (100 µL each). From each well, 10 µL were taken and serial dilutions were plated on LB agar plates (LB and 1.5 % agar; Carl Roth, Karlsruhe, Germany) to determine the number of colony forming units (CFUs) per mL in the inoculum. The 96-well plate was incubated at 37 °C for 4 h. Afterwards, a second serial dilution was performed to quantify the number of survived CFUs/mL. All LB agar plates were incubated overnight at 37 °C and formed colonies were counted the next day.

#### 2.3.4. Siderophore production

The ability to produce siderophores was tested as described elsewhere [38]. Briefly, colonies were suspended in 0.9 % (*w/v*) NaCl until OD_600_ reached 0.5 McFarland standard turbidity and diluted 1:100 in 5 mL iron-chelated M9 medium (200 μM 2,2′-dipyridyl [Carl Roth, Karlsruhe, Germany] added to M9 medium) supplemented with 0.3% (*w*/*v*) casamino acids (BD, Franklin Lakes, NJ, USA) in sterile 15 mL tubes (Sarstedt, Nürnbrecht, Germany). Following overnight incubation at 37°C and 130 rpm on a rotary shaker, 1 mL of the suspension was placed in 1.5 mL tubes and centrifuged at 4,900 × *g* and RT for 20 min. In a 96-well plate, 100 µL of the supernatant were mixed with 100 µL chrome azurol S shuttle solution (prepared according to [38]), 15 mM aqueous EDTA solution (Carl Roth, Karlsruhe, Germany) mixed with the shuttle solution was used as positive control, and the plate was incubated in the dark for 30 min at RT. The absorption was measured at λ = 630 nm with the microplate reader and siderophore production was calculated as previously described [39] and expressed as percentage unit of siderophore production.

### 2.4. Infection models

Larvae of the greater wax moth *Galleria mellonella* were infected with the isolates to test mortality as described previously [40]. Briefly, single colonies were suspended in PBS until an OD_600_ of 1 (approx. 2 × 10^9^ CFU/mL) was reached. After centrifugation at 12,000 × *g* and RT for 5 min, the bacterial suspensions were washed twice with PBS and diluted to 2 × 10^7^ CFU/mL. Then, larvae were randomly divided into groups of 10 individuals and 10 µL of the adjusted suspension were injected into the left proleg.

Additionally, 10 µL of PBS were injected into another group of larvae (n = 10) to exclude death due to injection trauma. Each group was placed into 90 mm Petri dishes and kept in the dark at 37 °C. Death was recorded every 24 h and larvae were considered dead when they no longer responded to physical stimuli and showed pigmentation. Results for the replicates of each isolate were pooled and Kaplan-Meier plots were used to show mortality rates [41].

*In vitro* adhesion was tested with the human epithelial cell lines 5637 and H3122, originating from urinary and pulmonal cancer, respectively. Cell lines were cultivated as previously described [42]. Cell culture medium for 5637 consisted of RMPI medium (Biowest, Nuaillé, France) supplemented with 10 % fetal bovine serum (Bio & Sell, Feucht, Germany), 2 mM glutamine (Biowest, Nuaillé, France) and 100 IU/100 µg per mL penicillin/streptomycin (Biowest, Nuaillé, France), while DMEM F12 medium (Biowest, Nuaillé, France) with the same supplements was used for H3122. Cell lines were grown at 37 °C with 5 % CO_2_ in cell culture flasks and passaged according to standard protocols. For adhesion assays, cell lines were seeded in 96-well plates (Nunclon, Thermo Fisher Scientific, Waltham, MA, USA) and grown to near confluence. Cells were then washed with 100 µL PBS and 100 µL of the corresponding cell culture medium without antibiotics were added. From blood agar plates, bacterial colonies were resuspended in the respective cell culture medium, then 50 µL of this suspension were added to the 96-well plate to match a multiplicity of infection of 100 (meaning 100 bacteria per cell). Six wells per plate were not inoculated with bacteria to serve as negative controls. Plates were incubated for 4 h at 37 °C with 5 % CO_2_ allowing bacteria to adhere to the cells. Then, the wells were washed thrice with 100 µL PBS and cells/bacteria were fixed with 4 % paraformaldehyde (in PBS) overnight at 4 °C. Afterwards, wells were washed thrice with 100 µL PBS and dehydrated with 50 µL 95 % ethanol for 5 min.

Fluorescence *in situ* hybridization (FISH) was used to stain the bacteria. The FISH probe Eub338 Atto647N (final concentration 5 ng/µL) was added to freshly prepared hybridization buffer (0.9 M NaCl, 20 mM TRIS-HCl, 0.01 % SDS, 15 % formamide) and 40 µL were given to each well. After incubation in a humidity chamber for 1 h at 46 °C, wells were washed with 100 µL prewarmed washing buffer (0.9 M NaCl, 20 mM TRIS-HCl, 0.01 % SDS) and incubated for 10 min at 48 °C, before washing once with 100 µL PBS. Nuclei were stained with 50 µL DAPI (50 µg/mL in H_2_O) and wells were washed thrice with 100 µL PBS. After washing, 100 µL PBS were left in each well. The plates were analyzed using the AKLIDES system (Medipan, Dahlewitz, Germany) and 10 images were taken from each well. Images were analyzed with MaxiSlider v.4.1.0 (Brandenburg University of Technology, Senftenberg, Germany), which determined the number of bacteria per mm^2^. Each image was manually checked for errors, e.g., insufficient number of human cells or high background noise, and potentially excluded from further analysis. The median adhesion for each isolate was calculated, blank corrected by subtracting the median of the negative control and averaged from all replications.

### 2.5. Microscopy

#### 2.5.1. Capsule staining

To identify the capsule, isolates 5A-D were used for capsule staining. Isolates were grown overnight on blood agar plates and each isolate was stained using the modified protocol of Maneval’s [43] method by Hughes and Smith (https://asm.org/Protocols/Capsule-Stain-Protocols). Here, a colony taken from the agar plate was mixed with 30 µL of 1 % Congo red initially pipetted onto a clean glass slide. The mixture was smeared using another slide and left to dry at RT. The air-dried slide was positioned on the rack of the staining tray. It was gently flooded with Maneval’s solution and left for 5 min. Subsequently, the slide was lifted, and the excess stain was carefully poured off. It was then placed inclinedly on the staining tray and rinsed gently with water. Finally, the slide was put on absorbent paper and left to dry at RT. Bright-field microscopy was performed using the Axio Imager.Z2m system equipped with the Plan-APOCHROMAT 100x/1.4 oil immersion objective (Carl Zeiss Microscopy, Oberkochen, Germany). Images were captured with the camera Axiocam 305 color using the ZEN 3.1 pro software (Carl Zeiss Microscopy, Oberkochen, Germany).

#### 2.5.2. Scanning electron microscopy

For further analysis of the different morphologies, scanning electron microscopy was performed for isolates 5A-D. Isolates were cultivated overnight on blood agar plates and, for each morphotype, colonies were resuspended in 1 mL 0.9 % NaCl until an OD_600_ of 1 was reached. This suspension was diluted with 9 mL of 0.9 % NaCl and filtered through a hydrophilic polycarbonate filter (pore size 0.2 µm; Merck Millipore, Darmstadt, Germany). A segment from the center of the filter was fixed with 1 % glutaraldehyde and 4 % paraformaldehyde in washing buffer (10 mM cacodylate buffer, 1 mM CaCl_2_, 0.075 % ruthenium red; pH 7) and samples were then stored at 4 °C until further processing. Samples were washed thrice with washing buffer for 10 min each, treated with 2 % tannic acid in washing buffer for 1 h at RT, and washed again thrice with washing buffer for 15 min each. Then, samples were dehydrated in a graded series of aqueous ethanol solutions (10 %, 30 %, 50 %, 70 %, 90 %, 100 %) on ice for 15 min for each step. Before the final change to 100 % ethanol, samples were allowed to reach RT and then critical point dried with liquid CO_2_. Finally, samples were mounted on aluminum stubs, sputtered with gold/palladium and examined with a field emission scanning electron microscope Supra 40VP (Carl Zeiss Microscopy, Oberkochen, Germany) using the Everhart-Thornley SE detector and the in-lens detector in a 70:30 ratio at an acceleration voltage of 5 kV. All micrographs were edited by using Adobe Photoshop CS6.

### 2.6. Data visualization and statistical analyses

For statistical analyses, GraphPad Prism v.7.05 (GraphPad software, San Diego, CA, USA) was used. Phenotypic experiments were performed with biological triplicates and each biological replicate was divided into technical triplicates. Due to higher variability, the CV assay was performed with six biological replicates, again divided into technical triplicates, and a ROUT outlier test was performed. Unless indicated otherwise, data are expressed as mean values and standard error of the mean. Statistical significance was determined with the Welch’s test (unpaired t test; two-tailed; confidence level 95 %), comparing the morphotypes of each time point to the white (normal) phenotype of that given time point, or analysis of variation (ANOVA) with Dunnett’s multiple comparison *post hoc* test, comparing all isolates to a given reference. Statistical significance is indicated by *p* values of less than 0.05. Synteny plots were generated with pyGenomeViz (https://github.com/moshi4/pyGenomeViz) and figures were created with BioRender.

## 3. Results

### 3.1. Increased resistance and virulence indicate acquisition of relevant genes including the chromosomal insertion of a large hybrid plasmid upon in-host evolution

As described in the study of Doğan et al., all *K. pneumoniae* isolates belonged to ST147 with an O locus of O1/O2v1 and a capsule biosynthesis (KL) locus of KL64 for all isolates except 4D, which missed most of its K locus [11]. Interestingly, all isolates with the grey or g/d phenotype lacked at least one K locus gene (Supplementary table S1). All isolates harbored the insertion sequence IS*1R* (IS*1* family, 768 bp) within the K locus and, except for 4D, also in the region between the K and O loci (disrupting an acyltransferase gene). In addition, we identified IS*1R* within the O locus (inserted into *wbbM*) of isolates 1A and 1B. While isolates 1A and 1B only carried Col plasmids (most similar to MGM828 [93.51 % identity] and pHAD28 [92.37 %]) and the incompatibility group (Inc)FIB(pKPHS1) (95.54 % identity), isolate 2B harbored additional IncFIAHI1 (97.17 % identity), IncFII(pKP91) (88.28 % identity) and IncR (100 % identity) groups (Supplementary table S1). All other isolates were additionally positive for IncHI1B(pNDM-MAR) (99.83 % identity) and IncFIB(pQil) (100 % identity) (Supplementary table S1). Investigation of the closed genomes of isolates 5A-D showed that two of the plasmids carried two Inc groups each (IncFIB[pQil] + IncHI1B[pNDM-MAR]; IncFIA[HI1] + IncFII[pKP91]), whereas the remaining four plasmids harbored one Inc group each. Accordingly, isolates 1A+B and 2B had lower Kleborate resistance scores than the later isolates (scores of 0 vs. 2) as described previously [11]. The virulence scores demonstrated the same trend (scores of 1 vs. 4). We detected several antimicrobial resistance features including ESBL and carbapenemase genes. These genes were located on the IncFIB(pQil) + IncHI1B(pNDM-MAR) plasmid and accordingly detected in all isolates except 1A+B and 2B. The isolates with low virulence only carried the siderophore genes for yersiniabactin, while the other isolates were additionally positive for aerobactin. Using the VFDB to assign additional virulence factors, up to 85 genes were identified per isolate (Supplementary table S2). The most abundant genes were associated with metabolic factors (n = 29), adherence and biofilm formation (n = 24), as well as effector delivery systems (n = 11). Metabolic factors included iron acquisition and transport such as the siderophore systems enterobactin (*entABCEFS*, *fepABCDG*, *fes*) yersiniabactin (*fyuA*, *irp1*, *irp2*, *ybtAEPQSTUX*) and aerobactin (*iutA*, *iucABCD*). Apart from *iutA*, aerobactin genes were not detected in 1A+B and 2B, which matched the virulence scores. Adherence genes included fimbriae and pili, while effector delivery systems were associated with type VI secretion systems. In addition, for all isolates except 1A+B, 2B and 5A, the plasmid-borne gene *rmpA2* (regulator of mucoid phenotype), one of the biomarker genes of hvKp classification [44], was detected. Finally, heavy metal resistance (HMR)-associated features likely conferring resistance against arsenic (*arsBCR*) were detected for all isolates, whereas 2A, 3A+B, 4A-D and 5A-D carried genes for tellurium HMR (*terABCDEYZ*) in addition. According to the analysis of the closed genomes, tellurium HMR was again located on the IncFIB(pQil) + IncHI1B(pNDM-MAR) plasmid. Taken together, these results indicate that plasmid-based acquisitions of resistance and virulence genes occurred during infection, which also led to an in-host transition from classic to convergent isolates.

Interestingly, in-depth analysis of the closed genome of 5D suggested that the IncFIB + IncHI1B plasmid with a size of approximately 355 kbp integrated into the bacterial host’s chromosome (Fig. 2). The plasmid is flanked by *ltrA* genes (group II intron reverse transcriptase/maturase) and inserted between *ltrA* and *spoIVCA* (DNA recombinase/invertase). Via reverse splicing, group II introns are able to insert genomic information directly into the chromosome [45]. Compared to the closed plasmid sequence of the 5B reference, the insertion shows one smaller region, containing *bla*_OXA-48_ and *csgG* (curli production assembly/transport component), with the same reading direction, and two large, inverted regions, containing the above mentioned EBSL and carbapenemase genes as well as the aerobactin operon (Fig. 2). PCR analysis (Supplementary figure S1) confirmed that the respective “plasmid-based” genes were only present in the chromosome of isolate 5D while being absent in 5A-C. Taken together, this clearly indicates chromosomal plasmid integration in isolate 5D.

**Figure 2.**
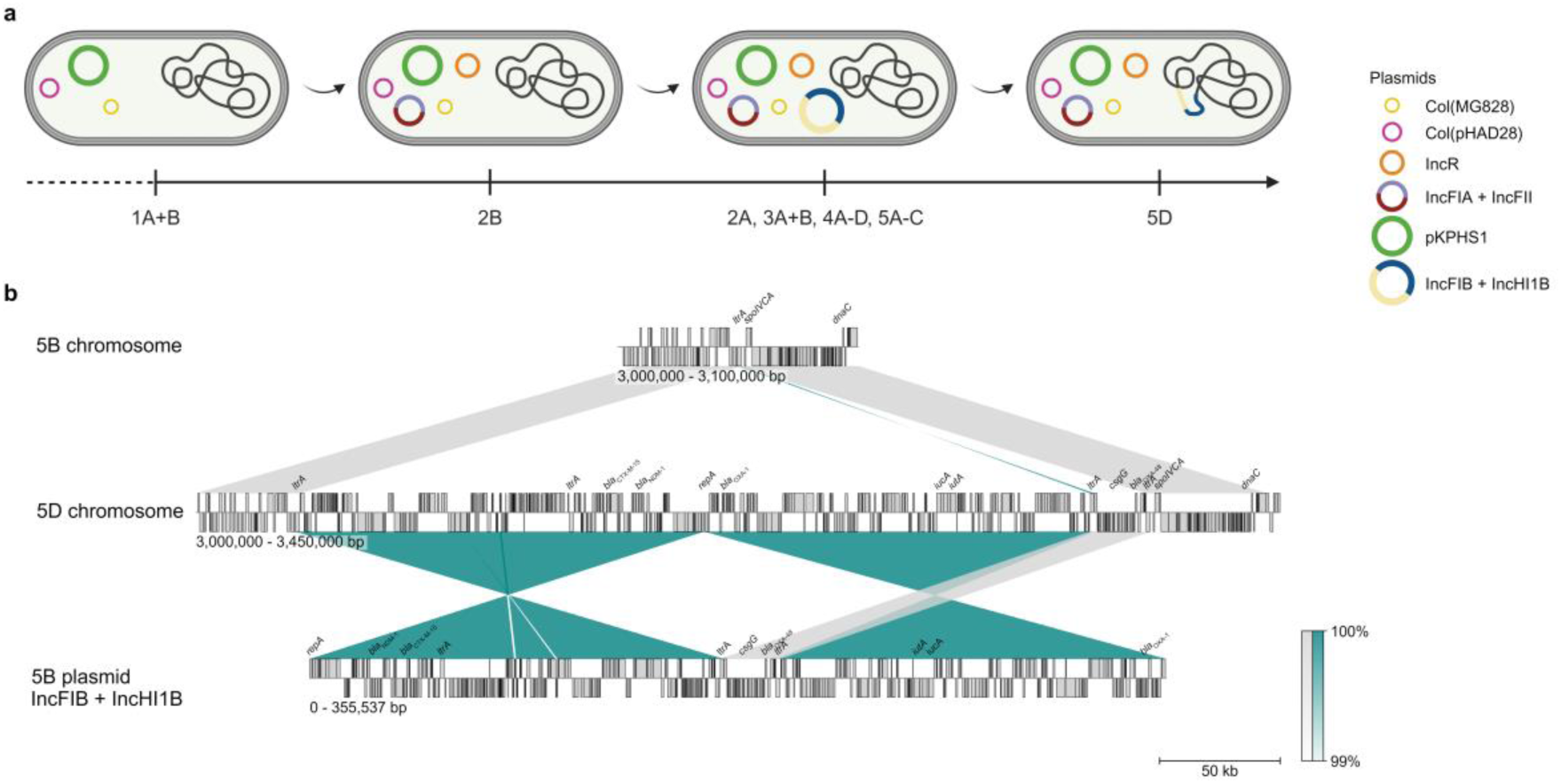
(**a**) Schematic representation of acquired plasmids over time. Despite sampled at different time points, the majority of the isolates carried the three initial and three acquired plasmids. One isolate showed a chromosomal integration of a large hybrid plasmid, carrying resistance and virulence genes. (**b**) Synteny plot of isolates 5B and 5D chromosomes and the IncFIB + IncHI1B plasmid. For clarity reasons, only selected resistance and virulence genes (e.g., *iutA* and *iucA*) are shown (*iucBCD* is excluded).

### 3.2. Variations in the K locus and adapted regulation of capsular genes likely contribute to the different morphotypes

When comparing all individual isolates, those obtained from the first three time points (1-3, both A+B) missed a 2,366 bp region within the chromosome including genes for a glycerol dehydrogenase and a transmembrane protein surrounded by IS*1* family transposases, that, in contrast, were present in isolate sets 4 and 5 (both A-D) indicating that this region was acquired later. Regarding the IncFIB + IncHI1B plasmid, it became apparent that isolates 4A and 5C lacked sequences of 777 bp and 3,954 bp in size, respectively. Both missing regions are likely linked to the IS*1R*, which is supported by the missing genes. In 4A, the region contained genes for a DUF4165-domain-containing protein and IS*1* family transposases. In 5C, the missing sequence harbored genes for a peptide deformylase (*def*), lysozyme inhibitor, transcriptional regulator and *cobW*, a C-terminal domain-containing protein, and was located between aerobactin-encoding genes and IS*1* family transposases.

Regarding the K locus, isolates 1A and 1B contained a missense mutation in *wzc* (c.1615C>A; p.P539T). All isolates, except 4D, 5C and 5D, carried a silent mutation in *wzy* (c.717A>G; p.V239V) and all isolates, except 4D, carried an additional missense mutation in *wcaJ* (c.305_306CG>AA; p.T102K). For isolates 2B and 3B, we identified a deletion within the K locus region mediated by IS*1R* and affecting genes *wzi*, *wza*, *wzb*, *wzc* (2B) and *wzi* and *wza* (3B), respectively. Further large deletions mediated by IS*1R* affected almost the complete K locus of isolate 4D (only *galF* and *cpsACP* present) and the genes *wzi*, *wza*, *wzb*, *wzc*, *wzx*, *wcoV*, a gene coding for a hypothetical protein and gene *wzy* of isolates 5C and 5D. In isolates 1B and 4C single K locus genes *wcoT* and *wcaJ*, respectively, were interrupted by IS*1R*. Notably, all isolates with the grey or g/d phenotype showed deletions in the K locus indicating that these morphotypes correlate with the K locus make up.

Next, we applied transcriptomics for a selected isolate set to further explore the underlying mechanisms of the different morphologies. We therefore compared the isolates 5A (small), 5C (grey) and 5D (g/d) to the white phenotype (5B), as this isolate demonstrated the closest morphology to “normal” *K. pneumoniae*. A total of 426 genes were differentially expressed in 5A (232 up- and 194 down-regulated), while we only found 23 (1 up- and 22 down-regulated) and 51 (28 up- and 23 down-regulated) genes in 5C and 5D, respectively (Supplementary figure S2). Regardless of the direction of regulation, shared differentially expressed genes among all three isolates, in 5A and 5C or in 5C and 5D, were chromosome-based and associated with the K locus, but mainly included undefined glycosyl transferases. Interestingly, these genes were up-regulated in the small colony variants (5A) compared to the white phenotype (5B), whereas they were down-regulated in the grey (5C) and g/d (5D) morphotypes. The down-regulation was partially because these genes were missing in 5C+D but present in the 5B reference (5/13 genes), whereas the remaining genes (8/13), e.g., *wcaJ*, were truly down-regulated. We could not assign shared differentially expressed genes of 5A and 5D to particular chromosomal regions, however, unannotated genes surrounding (and including) *csgG* were up-regulated in these isolates. We also detected unique genes regulated in only one morphotype. While 92.5 % of the regulated genes were unique to 5A (394/426, 204 up- and 190 down-regulated), only 21.7 % individually regulated genes were found in 5C (5/23, 1 up- and 4 down-regulated), and 35.3 % in 5D (18/51, 15 up- and 3 down-regulated). For 5C, the up-regulated gene belonged to an IS*1* family transposase, and the down-regulated genes were related to the missing region starting with *def* and ending with *cobW*. All unique 5D genes were chromosome-based and with some of the up-regulated genes belonging to multidrug efflux SMR transporters (*kpnEF*) or transferases (*argA*, *pspE*). The down-regulated genes were related to stress response (*pspG*) and sugar transport (*rbsB*). 5A presented up-regulated chromosomal genes from various groups, often related to DNA regulation, ABC transporters, ribosomal proteins, or transposases. Among the down-regulated genes were some of the enterobactin-encoding genes (*entABEF*, *fepCG*), conjugation (*traABDEIKLQSUV*, *trbCEFI*) and transposases. Interestingly, in contrast to the up-regulated chromosomal transposases, the down-regulated transposases were mainly plasmid-based. Overall, this indicates that differential expression of K locus genes contributes to the distinctive morphotypes.

### 3.3. Isolates from later time points demonstrate high virulence and certain morphotypes are associated with increased epithelial adhesion

To gain further insights into the association between the specific morphologies and bacterial virulence, fitness, and resilience, we performed several phenotypic assays and included two references, i.e., an archetypical hypervirulent (hvKP1) and a convergent *K. pneumoniae* strain (PBIO1953).

Note that only hvKP1 exhibited a positive string test and thus clear HMV, while in the sedimentation assay, in which mucus-producing cells sediment slower during the centrifugation process, isolate 1A produced significantly more mucus than hvKP1 (*p* = 0.0019) (Fig. 3a). When comparing the different morphotypes of each time point, isolates 1A and 4D showed higher sedimentation ratios than the corresponding isolates 1B (*p* = 0.0007) and 4B (*p* < 0.0001) (Fig. 3a). However, as none of the isolates demonstrated a positive string test result, they were not defined as hypermucoviscous. Siderophore production, which is an important indicator for bacterial virulence, was significantly lower for isolate 1A (*p* < 0.0001), 1B (*p* < 0.0001), 2B (*p* < 0.0001) and 4D (*p* = 0.0231) when compared to hvKP1 (Fig. 3b). Also note that isolate 2B produced significantly less siderophores than isolate 2A (*p* = 0.0168) (Fig. 3b). For 1A+B and 2B, this matched the WGS data, as these isolates missed genes for aerobactin, the dominant siderophore in *Klebsiella* [46]. The slightly lower siderophore production of 5A might be related to the down-regulation of some enterobactin-encoding genes as shown on transcriptomic levels. Interestingly, when challenged in 50 % human serum, all isolates with the grey and the g/d phenotypes had significantly lower CFU levels than hvKP1 (1B: *p* < 0.0001; 2B: *p* = 0.0041; 3B: *p* = 0.0417; 4C: *p* = 0.0045; 4D: *p* = 0.0006; 5C: *p* = 0.0004; 5D: *p* = 0.0004) after 4 hours of incubation (Fig. 3c). Isolates 3A and 4A were the only non-grey isolates with lower CFU values, however, these results were not significant. As *Klebsiella* often forms biofilms that are associated with intestinal colonization and subsequent infection, we investigated biofilm production properties in a crystal violet experiment. Biofilm production of hvKP1 was low and, with the exception of isolates 1A+B, 4D and PBIO1953, all isolates had significantly higher values (2A: *p* = 0.0002; 2B: *p* = 0.0029; 3A: *p* = 0.0144; 3B: *p* = 0.0125; 4A: *p* = 0.0378; 4B: *p* = 0.0258; 4C: *p* = 0.0326; 5A: *p* = 0.0363; 5B: *p* = 0.0304; 5C: *p* = 0.0121; 5D: *p* = 0.0304; W3110: *p* < 0.0001) (Fig. 3d). Compared to 4B, isolate 4D had a significantly lower specific biofilm factor (*p* = 0.0009) (Fig. 3d). Considering the higher mucosity of isolates 1A, 4D and hvKP1 suggests that biofilm formation might counteract mucus production. Despite the low biofilm production of some isolates, all grey and g/d phenotypes seemingly adhered better to human epithelial cells (Fig. 3e-f). Regarding the epithelial bladder cells (5637), only two isolates demonstrated significantly higher adhesion than hvKP1 (2B: *p* = 0.0098; 4D: *p* < 0.0001), while for the epithelial lung cells (H3122), several isolates attached significantly better (1B: *p* < 0.0001; 4C: *p* = 0.0043; 4D: *p* < 0.0001; 5C: *p* = 0.0014). In addition, isolates 1B and 4D showed significantly higher adherence than their associated, white “partners” (1B: *p* = 0.0053; 4D: *p* = 0.0196). In general, most isolates displayed higher attachment to epithelial lung (H3122) than to bladder cells (5637).

**Figure 3.**
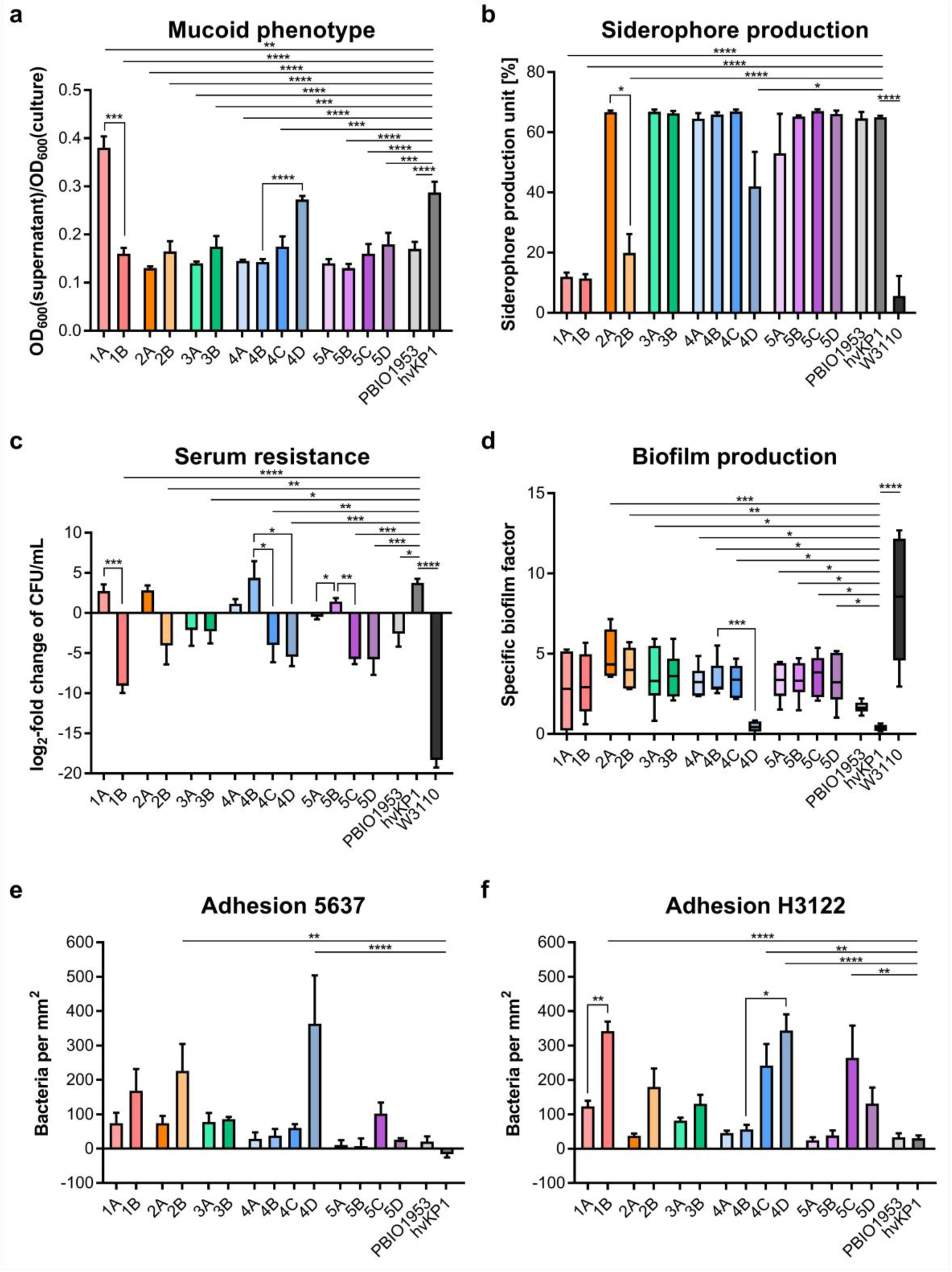
Phenotypic assays to determine virulence and resilience of *K. pneumoniae*. (**a**) Determination of the mucoid phenotype using a sedimentation assay. Results are given as mean ratios of OD_600_ of supernatant after centrifugation and total OD_600_ and standard error of the mean (n = 3). (**b**) Siderophore production of the respective isolates, shown as mean siderophore production unit and standard error of the mean (n = 3). (**c**) Survival in 50 % human serum after 4 hours of incubation. Results are given as mean log_2_-fold change of CFU/mL and standard error of the mean (n = 3). (**d**) Boxplots of biofilm production given as the specific biofilm factor. Mean is indicated within the boxes, whiskers show 10-90 percentile of standard error of the mean (n = 6). (**e**) Adhesion of the respective isolates to 5637 epithelial bladder cells. Results are given as mean number of attached bacteria per mm2 and standard error of the mean (n = 3). (**f**) Adhesion of the respective isolates to H3122 epithelial lung cells. Results are given as mean number of attached bacteria per mm2 and standard error of the mean (n = 3). Statistical significance was tested with one-way ANOVA (with Dunnett’s multiple comparison post hoc test), indicated by the straight bars, and with Welch’s t test, indicated by the brackets; * p < 0.05, ** p < 0.01, *** p < 0.001, **** p < 0.0001.

Finally, we explored the isolates’ ultimate virulence potential in an *in vivo* mortality assay using *Galleria mellonella* larvae (Fig. 4). Surprisingly, many isolates had higher mortality rates than the hvKP1 reference, however, the differences were not significant. Two of the three isolates with the g/d phenotype (1B and 4D) exhibited significantly lower mortalities for all three analysis time points (24h, 48h and 72h) (Fig. 4). In addition, the two isolates with the small colony morphologies (4A and 5A) had rather low mortalities with 5A differing significantly from hvKP1 for the last two analysis time points. Overall, most of the isolates, especially the later ones, demonstrated high virulence and resilience, but we did not detect a clear association with specific phenotypes.

**Figure 4.**
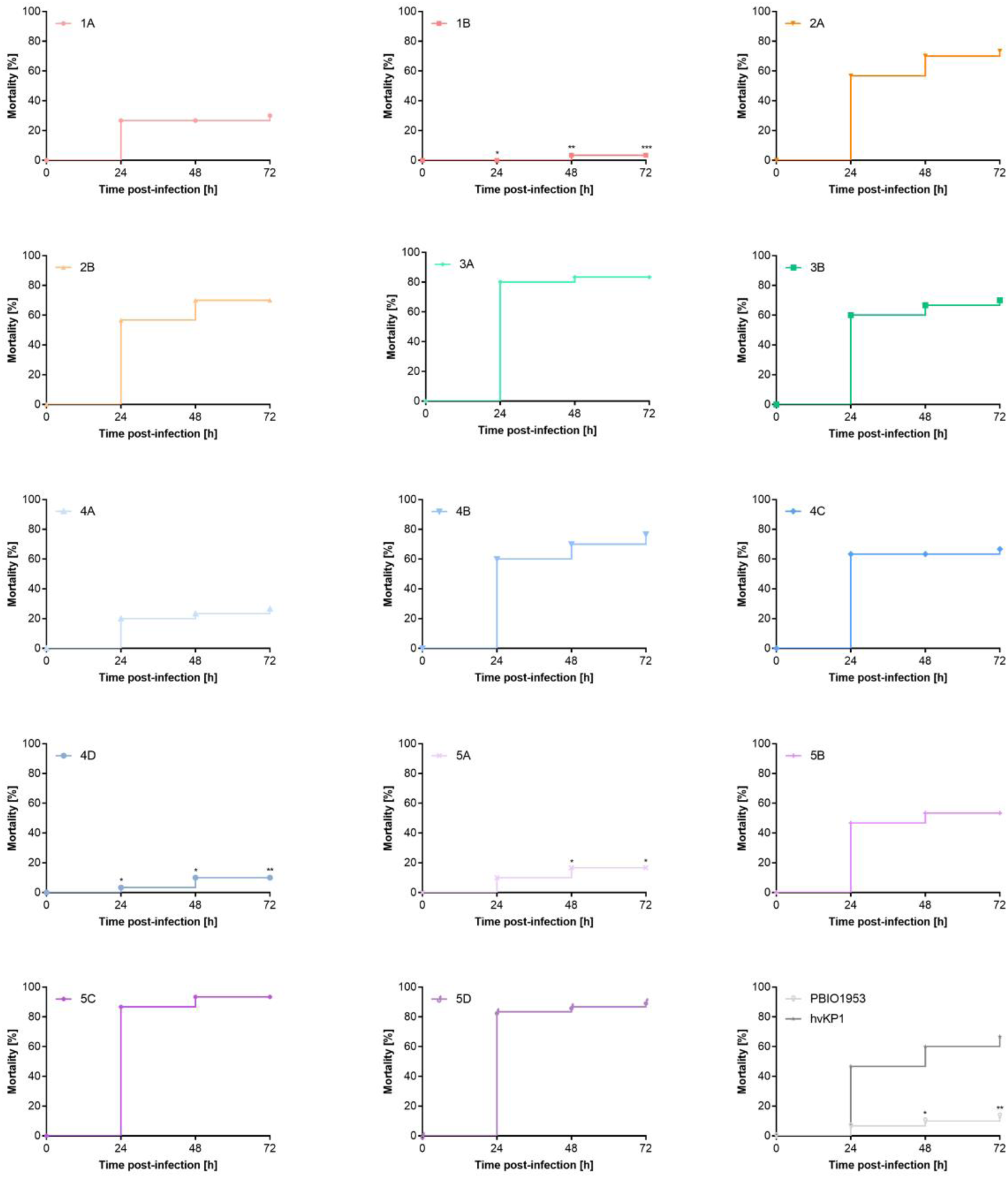
Kaplan-Meier plots of mortality rates of *Galleria mellonella* larvae in the infection model. Results are given as mean mortality after injection of 2 × 10^5^ CFU. Statistical significance was tested with two-way ANOVA (with Dunnett’s multiple comparison post hoc test); * p < 0.05, ** p < 0.01, *** p < 0.001.

### 3.4. Microscopy results indicate altered cell size and decreased capsule production of certain morphotypes

While the capsule staining proved encapsulations of all isolates (5A-D), differences were detected regarding their size or distribution (Fig. 5.1). The small and the white morphotypes showed some clustering (Fig. 5a.1, 5b.1), while the grey and g/d phenotypes displayed a more even distribution (Fig. 5c.1, 5d.1). Interestingly, several elongated cells were visible for the g/d phenotype (Fig. 5d.1).

**Figure 5.**
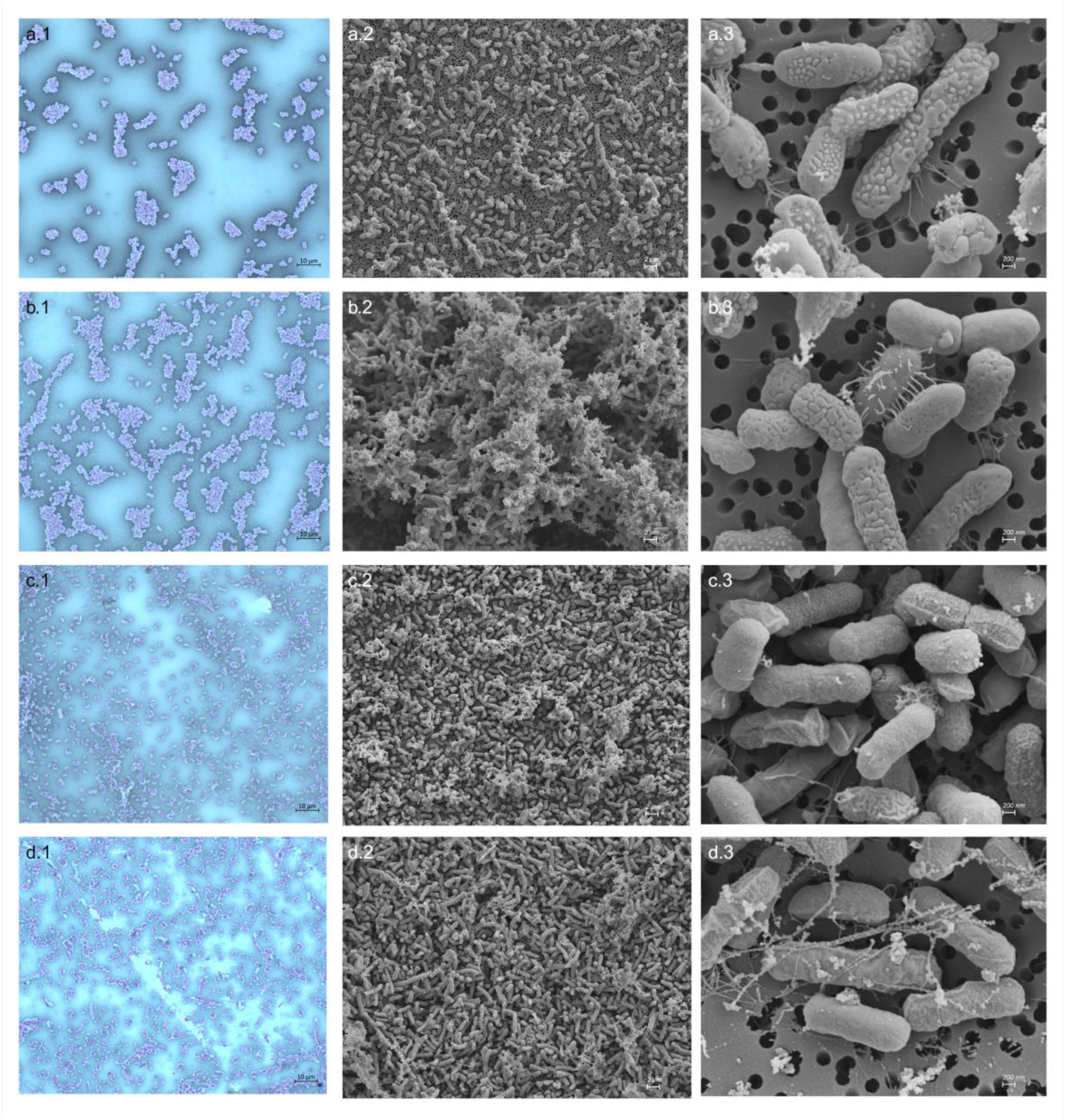
Micrographs of the (**a**) small (5A), (**b**) white (5B), (**c**) grey (5C) and (**d**) grey and dry (5D) phenotype. Left column: Capsule stain using the modified Maneval’s method by Hughes and Smith, scale bars = 10 µm. Middle column: Scanning electron micrographs at 2,000x magnification, scale bars = 2 µm. Right column: Scanning electron micrographs at 20,000x magnification, scale bars = 200 nm.

To visualize the different morphotypes in high-resolution, we also performed scanning electron microscopy of the isolates 5A-D (Fig. 5a.2-5d.2, 5a.3-5d.3). Whereas the hypervirulence reference strain PBIO2030 had rather small cells with some inter-cell linkages (Supplementary figure 3b) and a thick capsule (Supplementary figure 3c), the small phenotype (5A) presented a variety of differently sized cells and only few connections between cells (Fig. 5a.2). Here, the capsules revealed inconsistent structures, some areas on the same cell appeared similar as PBIO2030, but others demonstrated a more uniform pattern, possibly indicating that some surface areas contain more capsular polysaccharides than others (Fig. 5a.3). The white phenotype (5B) presented mostly medium-sized highly aggregated cells (Fig. 5b.2) and with a less disrupted capsule, pointing towards the production of a “normal” amount of capsular polysaccharides (Fig. 5b.3). Similar applied to the grey phenotype (5C) but with less capsule production (Fig. 5c.1, 5c.2). Isolate 5D (g/d) displayed a high number of elongated cells, as previously seen during the capsule staining, with many interconnections (Fig. 5d.2) and a similar capsule as 5C, indicating lower capsular polysaccharide production. While the fine-structured linkages that are directly connected to cells, as seen for 5B (Fig. 5b.3), are probably fimbrial structures, the longer and cruder looking linkages, especially seen for 5D (Fig. 5d.3), are extracellular polysaccharides, e.g., biofilm components. As isolates 5C and 5D exhibited an identical K locus make up, their different morphologies might be related to cell size and inter-cell linkages, related to biofilm production, which might be associated with the upregulation of the *csgG* region for 5D. Taken together, the grey (5C) and g/d (5D) morphotypes seem to produce lower amounts of capsular polysaccharides, which correlates with the K locus make up and the adapted regulation of the corresponding genes.

## 4. Discussion

The isolates characterized in this study belong to *K. pneumoniae* ST147, an international high-risk clonal lineage [47,48]. Using Kleborate, Lam et al. investigated several *K. pneumoniae* genomes, including 271 genomes of ST147 [47]. The most common resistance score among ST147 genomes was 2, which applied to 52.8 % (143/271), and the least common score was 0, reflecting 8.5 % (23/271) [47]. This matches our results, where most isolates had a resistance score of 2. Also, in this other study, virulence scores of 0 (50.6 %, 137/271) and 1 (43.9 %, 119/271) were frequent, yet a virulence score of 4 was rare (1.1 %, 3/271) [47], which was the virulence score we detected the most (78.6 %, 11/14). Whereas the first isolates (1A+B, 2B) had low virulence scores, the resistance and virulence scores (2 and 4, respectively) of the later isolates (2A, 3A+B, 4A-D, 5A-D) suggest in-host evolution and a “transition” from the classic to the convergent pathotype. This was mainly due to the acquisition of additional plasmids, that likely originate from other bacteria within the host or within the clinical setting. The convergent character was confirmed on phenotypic levels. While cases of convergent *K. pneumoniae* are becoming more frequent [49,50], to the best of our knowledge, this is the first study demonstrating its *in vivo* evolution in a patient. Convergent strains are mainly a result of either AMR plasmid acquisition by hvKp, hypervirulence plasmid acquisition by multidrug-resistant cKp or the uptake of hybrid plasmids [50,51]. The latter is seemingly the case for our later isolates, which acquired the IncFIB+IncHI1B hybrid plasmid. Even more interesting, our analysis confirmed chromosomal integration of this plasmid in isolate 5D, potentially leading to fixation of the beneficial virulence and resistance features [10], stable propagation [52] and possibly decreased fitness costs compared to isolates that carry the respective genes on plasmids [21]. Like in our case, chromosomal plasmid insertion related to *ltrA* genes has been previously described, e.g., for *Clostridium* spp., where both the original plasmid and chromosomal target region contained several *ltrA* introns in close proximity to the integration site [53].

All our grey or g/d morphotypes missed genes within the K locus. While isolates 1B and 4C lacked only one gene, 3B missed two genes and 2B four. Isolate 5C and 5D each lacked eight genes and 4D missed almost all K locus genes. This accorded to high numbers of IS elements within and around the K loci that are likely responsible for the deletions. That is, identical IS elements within the same genome often drive genomic rearrangements, including deletion events [54]. Note that the hyper-splitting phenomenon is also related to the impaired K loci due to deleterious recombinations, which are responsible for the irreversible switch from small to grey or g/d phenotypes. The up-regulation of several DNA regulators and the large number of differentially expressed genes in general might indicate that the small phenotype is related to transcriptomic adaptations rather than adaptations on a genomic level.

As seen in the biofilm assay, the truncated K locus of 4D might lead to decreased adhesion to abiotic surfaces, since capsular polysaccharides play an important role in initial adhesion steps [55,56]. However, adhesion to biotic surfaces was not impaired. In fact, all isolates with the grey or g/d phenotype showed increased attachment to epithelial cells, which might be a result of differentially regulated K locus genes. Also, despite that the grey and g/d morphotypes demonstrated better attachment to epithelial cells, this simultaneously led to decreased serum resistance. As suggested by capsule staining and electron microscopy, these isolates produced less amounts of capsule, potentially resulting in a higher bacterial vulnerability to immune responses [57]. Furthermore, the relatively low serum resilience of isolates 4A and 5A might be due to their small-colony morphologies, as SCVs are usually associated with intracellular persistence [12,58] and thus only little exposure to the host’s immune system. The potentially invasive nature of the small phenotypes and their possible adaptation to an intracellular lifestyle could explain the rather low lethality caused by these isolates in the *G. mellonella* larvae. We speculate that, instead of inducing acute infection that results in larvae death, the small isolates persist within the cells as it was demonstrated for staphylococcal SCVs [59]. It is noteworthy, that the intracellular lifestyle of staphylococcal SCVs has been associated with chronic persistent and recurrent courses of infection that are difficult to detect and treat [13,59]. The small *K. pneumoniae* phenotype isolates in this study displayed a wide range in the size of its cells and few intercellular connections. Also for staphylococcal SCVs, heterogenous cells of different sizes have been seen in scanning electron microscopy [60].

Interestingly, Ernst et al. described clinical *K. pneumoniae* ST258 isolates presenting as grey, flat colonies on blood agar [7] that were determined “hypomucoid” and very similar to our g/d morphotypes. While these isolates were particularly associated with urinary tract infection [7] and bacterial capsule deficiency, “hypermucoid” representatives from the same study caused bloodstream infections and produced a hypercapsule [7]. Like in our isolates, the authors documented large deletions and/or IS elements within the K locus of the grey types [7]. However, their isolates with impaired K loci exhibited increased persistence but unchanged virulence or dissemination [7], but the latter was increased for our isolates due to the acquisition of virulence-associated genes. We agree that a thick capsule is considered an essential virulence factor in bacteria because of, e.g., mediation of phagocytosis protection [57]. However, capsule-deficient isolates potentially also lead to severe infection and should not be overlooked regarding their clinical relevance.

### 4.1. Conclusion

In conclusion, our study tackled the in-host evolution from classic *K. pneumoniae* to the convergent pathotype and revealed the chromosomal integration of a large hybrid plasmid for one isolate. We also demonstrated that genomic alterations, i.e., deletions within the K locus, caused the grey and g/d morphotypes that were also related to an increased adhesion to epithelial cells, and that the small phenotype was rather based on transcriptomic adaptations.

## Data availability statement

The data for this study has been deposited in the European Nucleotide Archive (ENA) at EMBL-EBI under the accession number PRJEB71325 (https://www.ebi.ac.uk/ena/browser/view/PRJEB71325).

## Ethics statement

Ethical review and approval were waived for this study, since this manuscript contains no personal information.

## Acknowledgement

The authors thank Stefan Bock for the excellent technical assistance regarding electron microscopy.

## Authors contributions

K.B. and K.Sc. conceived and designed the study. K.Sy. performed the laboratory and phenotypic experiments. E.A.I., E.D., E.E. and R.S. performed the microscopic experiments. M.S. and S.E.H. performed the bioinformatics analyses. D.B., E.A.I., E.D., E.E., J.A.B., J.U.M., K.B., K.Sy., K.Sc., M.M.K., M.S., P.S., R.S., and S.E.H. analyzed and interpreted the results. K.Sc. and K.Sy. drafted the manuscript with input from all co-authors, and K.Sy. and M.S. visualized the results. All authors read and approved the final version of the manuscript.

## Appendix. Supplementary data

Supplementary table S1. Results of whole-genome sequencing concerning sequence type, virulence and resistance scores, K and O loci and plasmids.

Supplementary table S2. Detected virulence factors according to VFDB.

**Supplementary figure S1.**
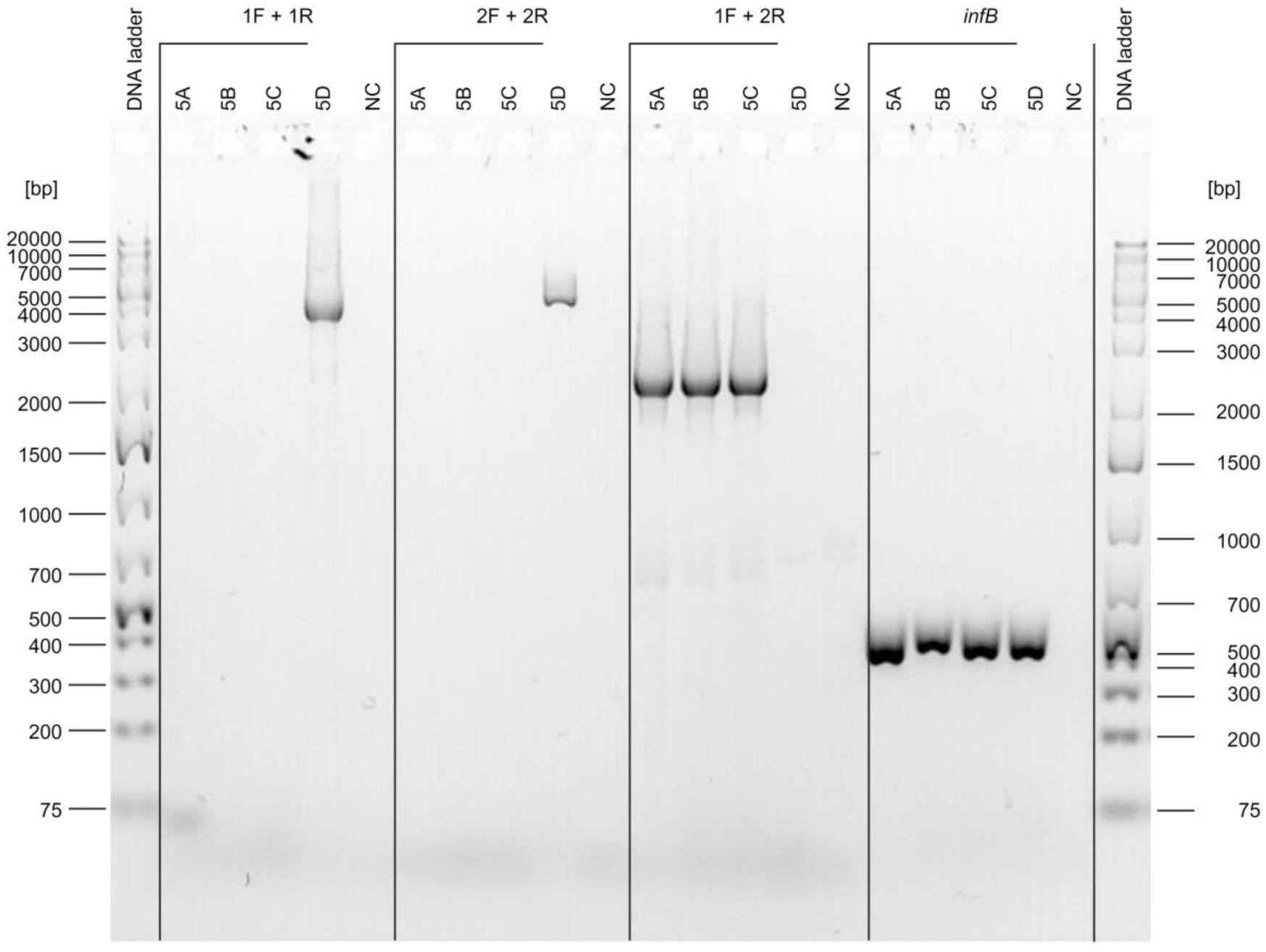
Representative gel electrophoresis image of PCR products supporting the chromosomal plasmid integration of isolate 5D. Specific primer pairs were used for the amplification of the left and right junction sequences of the chromosome and integrated plasmid. The expected amplicons were sized 4,499 bp (primer 1F + 1R, left junction) and 4,882 bp (2F + 2R, right junction), respectively. Additionally, the primers 1F and 2R were combined to exclude that no insertion was present in isolate 5D and confirm there was no insertion for the other isolates (5A-C). Genomic DNA was used as template and the negative controls (NC) were substituted with water instead.

**Supplementary figure S2.**
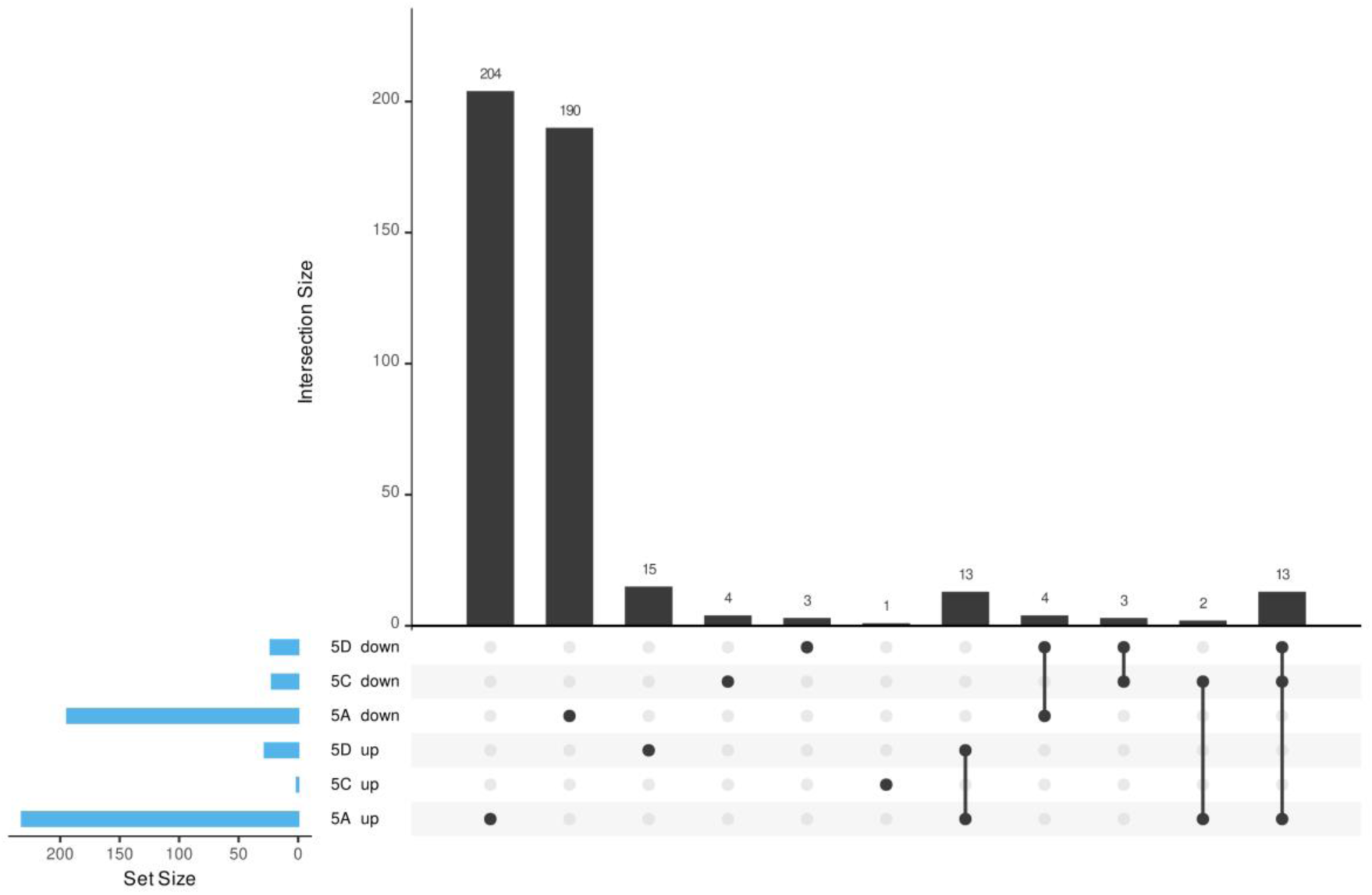
UpSet plot of differentially expressed genes in the different phenotypes. All phenotypes (5A: small; 5B: grey; 5C: grey and dry) were compared to the white phenotype (5B).

**Supplementary figure S3.**
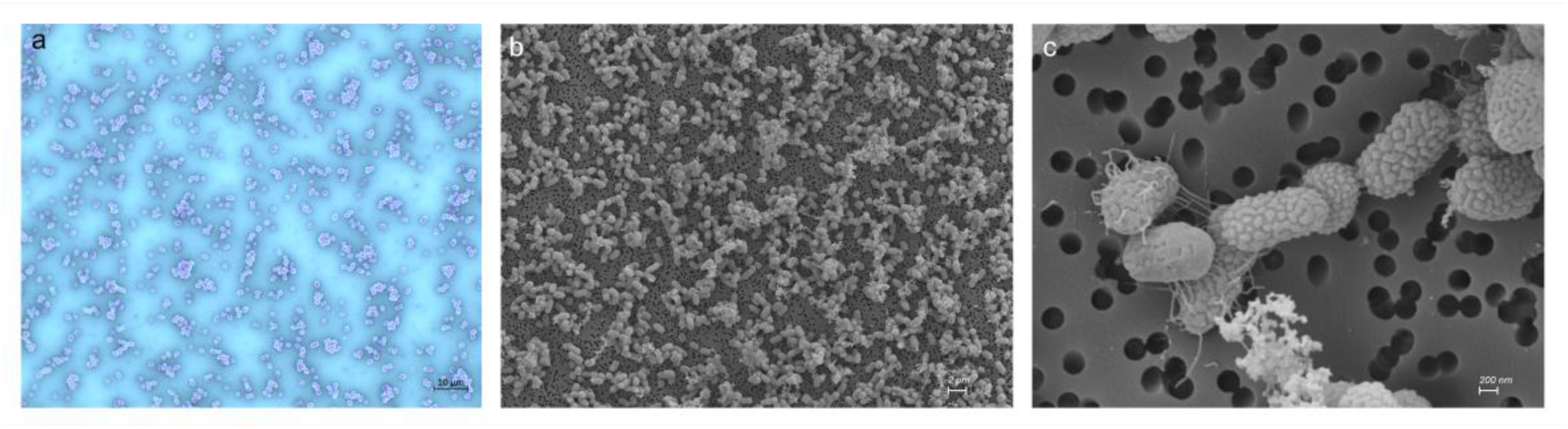
(**a**) Capsule stain of *K. pneumoniae* ATCC 13883 using the modified Maneval’s method by Hughes and Smith, scale bar = 10 µm. (**b**) Scanning electron micrograph of the hypervirulent *K. pneumoniae* PBIO2030 at 2,000x magnification, scale bar = 2 µm, and (**c**) at 20,000x magnification, scale bar = 200 nm.

